# An improved codon modeling approach for accurate estimation of the mutation bias

**DOI:** 10.1101/2021.06.30.450338

**Authors:** T. Latrille, N. Lartillot

**Affiliations:** Université de Lyon, Université Lyon 1, CNRS, Laboratoire de Biométrie et Biologie Évolutive UMR 5558, F-69622 Villeurbanne, France; École Normale Supérieure de Lyon, Université de Lyon, Université Lyon 1, Lyon, France

**Keywords:** codon models, phylogenetics, nucleotide bias, mutation-selection models

## Abstract

Nucleotide composition in protein-coding sequences is the result of the equilibrium between mutation and selection. In particular, the nucleotide composition differs between the three coding positions, with the third position showing more extreme composition than the first and the second positions. Yet, phylogenetic codon models do not correctly capture this phenomenon and instead predict that the nucleotide composition should be the same for all 3 positions of the codons. Alternatively, some models allow for different nucleotide rates at the three positions, a problematic approach since the mutation process should in principle be blind to the coding structure and homogeneous across coding positions. Practically, this misconception could have important consequences in modelling the impact of GC-biased gene conversion (gBGC) on the evolution of protein-coding sequences, a factor which requires mutation and fixation biases to be carefully disentangled. Conceptually, the problem comes from the fact that phylogenetic codon models cannot correctly capture the fixation bias acting against the mutational pressure at the mutation-selection equilibrium. To address this problem, we present an improved codon modeling approach where the fixation rate is not seen as a scalar anymore, but as a tensor unfolding along multiple directions, which gives an accurate representation of how mutation and selection oppose each other at equilibrium. Thanks to this, this modelling approach yields a reliable estimate of the mutational process, while disentangling fixation probabilities in different directions.

## 1 Introduction

Phylogenetic codon models are now routinely used in many domains of bioinformatics and molecular evolutionary studies. One of their main applications has been to characterize the genes, sites (Nielsen and Yang, 1998; Yang *et al*., 2005; Murrell *et al*., 2012) or lineages (Zhang and Nielsen, 2005; Kosakovsky Pond *et al*., 2011) having experienced positive selection (Murrell *et al*., 2015; Enard *et al*., 2016). More generally, these models highlight the respective contributions of mutation, selection, genetic drift (Teufel *et al*., 2018) and biased gene conversion (Pouyet and Gilbert, 2020; Kosiol and Anisimova, 2019), and the causes of their variation between genes (Zhang and Yang, 2015) or across species (Seo *et al*., 2004; Popadin *et al*., 2007; Lartillot and Poujol, 2011).

Conceptually, codon models take advantage of the fact that synonymous and non-synonymous substitutions are differentially impacted by selection. Assuming synonymous mutations are neutral, the synonymous substitution rate is equal to the underlying mutation rate (Kimura, 1983). Non-synonymous substitutions, on the other hand, reflect the combined effect of mutation and selection (Ohta, 1995). Classical codon models formalize this idea by invoking a single parameter *ω*, acting multiplicatively on non-synonymous substitutions rates (Muse and Gaut, 1994; Goldman and Yang, 1994). Using a parametric model automatically corrects for the multiplicity issues created by the complex structure of the genetic code and by uneven mutation rates between nucleotides. As a result, *ω* captures the net, or aggregate, effect of selection on non-synonymous mutations, also called *d*_*N*_ */d*_*S*_ (Spielman and Wilke, 2015; Dos Reis, 2015).

Classical codon models, so defined, are phenomenological, in the sense that they capture a complex mixture of selective effects through a single parameter (Rodrigue and Philippe, 2010). In reality, the selective effects associated with non-synonymous mutations depends on the context (site-specificity) and the amino acids involved in the transition (Kosiol *et al*., 2007). Attempts at an explicit modelling of these complex selective landscapes have also been done, leading to mechanistic codon models, based on the mutation-selection formalism (Halpern and Bruno, 1998). These models, further developed in multiple inference frameworks (Rodrigue *et al*., 2010; Tamuri and Goldstein, 2012), sometimes using empirically informed fitness landscapes (Bloom, 2014), could have many interesting applications, such as inferring the distribution of fitness effects (Tamuri and Goldstein, 2012) or detecting genes under adaptation (Rodrigue and Lartillot, 2016; Rodrigue *et al*., 2021), or even phylogenetic inference (Ren *et al*., 2005). However, they are computationally complex and potentially sensitive to the violation of their assumptions about the fitness landscape (such as site independence). For this reason, phenomenological codon models remain an attractive, potentially more robust, although still perfectible approach.

The parametric design of typical codon models, relying on a single aggregate parameter *ω*, raises the question whether they reliably estimate the underlying mutational process. Several observations suggest that this may not be the case. For instance, in their simplest form (Muse and Gaut, 1994; Goldman and Yang, 1994), codon models predict that the nucleotide composition should be the same for all three positions of the codons, and should be equal to the nucleotide equilibrium frequencies implied by the underlying nucleotide substitution rate matrix. In reality, the nucleotide composition differs: the third position shows more extreme GC composition, reflecting the underlying mutation bias, compared to the first and second positions, which are typically closer to 50% GC (Singer and Hickey, 2000).

These modulations across the three coding positions have been accommodated using the so-called 3×4 formalism (Goldman and Yang, 1994; Pond and Muse, 2005a), allowing for different nucleotide rate matrices at the three coding positions. However, this is also problematic, since this modelling approach has the consequence that synonymous substitutions, say, from A to C, occur at different rates at the first and third positions. Yet, in reality, the mutation process is blind to the coding structure, and should be homogeneous across coding positions, and if neutral, all mutations from A to C should thus have the same rate.

These observations suggest that the mutation matrix (1×4) or matrices (3×4) estimated by codon models are not correctly reflecting the mutation rates between nucleotides (Rodrigue *et al*., 2008; Kosakovsky Pond *et al*., 2010). Instead, what these matrices are capturing is the result of the compromise between mutation and selection at the level of the realized nucleotide frequencies. For detecting selection, this problem is probably minor, although it still bears consequences on the estimation of *ω* (Spielman and Wilke, 2015). Conceptually, however, it is a clear symptom of a more fundamental problem: mutation rates and fixation probabilities are not correctly teased apart by current codon models.

Practically, this misconception could have important consequences in contexts other than tests of positive selection. In particular, there is a current interest in investigating the variation between species in GC content, and its effect on the evolution of protein-coding sequences. An important factor here is biased gene conversion toward GC (called gBGC), which can confound the tests for detecting positive selection and, more generally, the estimation of *ω* (Galtier *et al*., 2009; Ratnakumar *et al*., 2010; Lartillot *et al*., 2013; Figuet *et al*., 2014; Bolívar *et al*., 2019). Even in the absence of gBGC, however, uneven mutation rates varying across species can have an important impact on the estimation of the strength of selection (Guéguen and Duret, 2018). All this suggests that, even before introducing gBGC in codon models, correctly formalizing the interplay between mutation and selection in current codon models would be an important first step.

In this direction, the key point that needs to be correctly formalized is the following. If the nucleotide’s realized frequencies are the result of a compromise between mutation and selection, then this implies that the strength of selection is not the same between all nucleotide or amino-acid pairs. For instance, if the mutation process is AT-biased, then, because of selection, the realized nucleotide frequencies at equilibrium will be less AT-biased than expected under the pure mutation process. However, this implies that, at equilibrium, there will be a net mutation pressure toward AT, which has to be compensated for by a net selection differential toward GC.

All this suggests that, in order for a codon model to correctly formalize this subtle interplay between mutation and selection, the component of the parameter vector responsible for absorbing the net effect of selection (i.e. *ω*) should not be a scalar, as is currently the case. Instead, it should be a tensor, that is, an array of *ω* values unfolding along multiple directions. In the present work, we address the question of whether we can derive a parametric structure being able correctly tease apart mutation rates and selection, and this, without having to explicitly model the underlying fitness landscape. In order to derive a codon model along those lines, our strategy is to first assume a true site-specific evolutionary process, following the mutation-selection formalism. Then, we derive the mean substitution process implied across all sites by this mechanistic model and identify the mean fixation probabilities appearing in this mean-field process with the *ω* tensor to be estimated. Inferring parameters on simulated alignments, we show that the model correctly estimates the mutation rates, as well as the mean effect of selection.

## 2 Results

To illustrate the problem, we first conduct simulation experiments under a simple mutation-selection substitution model assuming site-specific amino-acid preferences. We use these simulation experiments to explore through summary statistics the intricate interplay between mutation and selection. Then, we explore how codon models with different parameterizations are able to infer the mutation rates and the strength of selection on these simulated alignments. Finally, these alternative models are applied to empirical data.

### 2.1 Simulations experiments

Simulations of protein-coding DNA sequences were conducted under an origination-fixation substitution process (McCandlish and Stoltzfus, 2014) at the level of codons (see section 4.1). We assume a simple mutation process with a single parameter controlling the mutational bias toward AT, denoted *λ* = (*σ*_*A*_ + *σ*_*T*_)*/*(*σ*_*C*_ + *σ*_*G*_), where *σ*_*x*_ is the equilibrium frequency of nucleotide *x*. This mutational process is shared by all sites of the sequence. With regards to selection, synonymous mutations are considered neutral, such that the synonymous substitution rate equal to the underlying mutation rate. At the protein level, selection is modelled by introducing site-specific amino-acid fitness profiles (i.e. a vector of 20 fitnesses for each coding site), which are scaled by a relative effective population size *N*_r_. A high *N*_r_ induces site-specific profiles having a large variance, with some amino acids with a high scaled fitness while all other have a low scaled fitness. Conversely, a low value for *N*_r_ induces more even amino-acid fitness profiles (i.e. neutral) at each site. Thus, ultimately, the stringency of selection increase with *N*_r_. Altogether, the two parameters of the model tune the mutation bias (*λ*) and the stringency of selection (*N*_r_), respectively. All simulations presented are obtained using the same underlying tree topology and branch lengths of 61 primates from Perelman *et al*. (2011), and 4980 codon sites with amino-acid fitness profiles resampled from experimentally determined profiles in Bloom (2017).

Simulation of this origination-fixation process along a species tree result in a multiple sequence alignment of coding sequences for the extant species, from which summary statistics can then be computed. One such straightforward summary statistic is the frequency of the different nucleotides, and the resulting nucleotide bias AT/GC observed in the alignment. This observed nucleotide bias can be computed separately for each coding position (first, second and third) and compared to the underlying true mutational bias *λ*. As can be seen from figure 1, the third position of codons (panel C) reflects the underlying mutational bias quite faithfully, while the first and second positions (panel A and B) are impacted by the strength of selection and display nucleotide biases that are less extreme than the one implied by the mutational process. This differential effect across the three coding positions is explained by nucleotide mutations at the third codon position being more often synonymous, while mutations at the first and second positions are more often changing the amino-acid and are thus more often under purifying selection.

**Figure 1:**
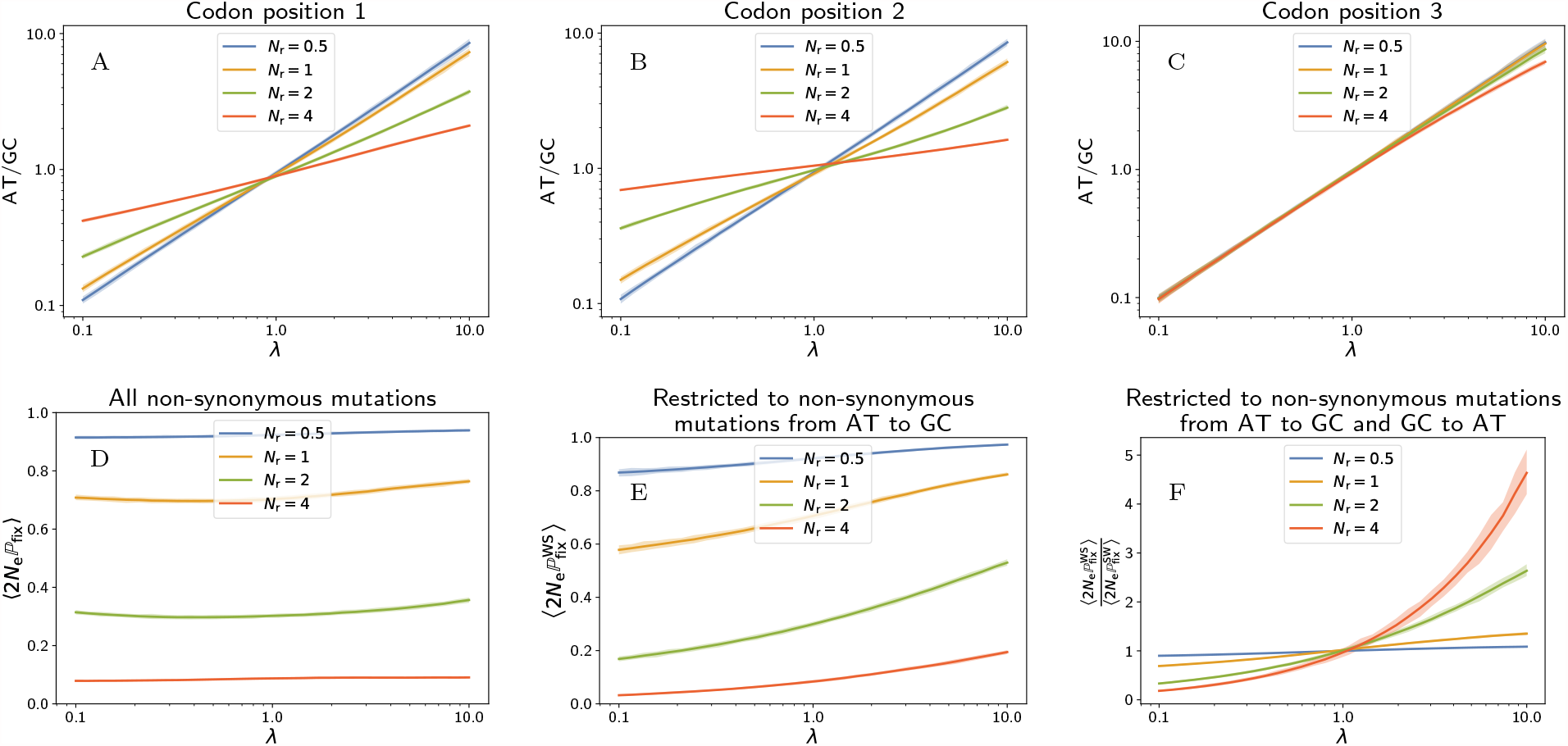
Simulations of 61 primates taxa, 4980 codon sites, with 100 repeats. Solid lines represent the mean value over the repeats, and the colored area the 95% inter-quantile range. Top row (A-C): Observed AT/GC composition of simulated alignment (first, second and third coding positions), as a function the underlying mutational bias towards AT (*λ*), under different stringencies of selection (different values of effective population size *N*_r_). Bottom row (D-E): Mean scaled fixation probability of non-synonymous mutations along simulations, ⟨2*N*_e_ℙ_fix_⟩, for all mutations (D) and for AT-to-GC mutations only (E), as a function of the mutational bias (*λ*), under different effective population sizes (*N*_r_). F: Ratio of mean scaled fixation probability for AT-to-GC over GC-to-AT mutations, as a function of the mutational bias and under different stringencies of selection (*N*_r_). Mutational bias is balanced by selection in the opposite direction, where this effect increases with the stringency of selection.

Apart from the observed nucleotide bias in the alignment, a statistic directly relevant for measuring the intrinsic effect of selection is the mean scaled fixation probability of non-synonymous mutations, called ⟨2*N*_e_ℙ_fix_⟩. This summary statistic ⟨2*N*_e_ℙ_fix_⟩ can be quantified from the substitutions recorded along the simulation trajectory (see section 4.4). For very long trajectories, it identifies with the ratio of non-synonymous over synonymous substitution rates (or *d*_*N*_ */d*_*S*_) induced by the underlying mutation-selection model (Spielman and Wilke, 2015; Dos Reis, 2015; Jones *et al*., 2017). As expected, ⟨2*N*_e_ℙ_fix_⟩ is always lower than 1 for simulations at equilibrium, under a time-independent fitness landscape (Spielman and Wilke, 2015). Quite expectedly ⟨2*N*_e_ℙ_fix_⟩ decreases with the *N*_r_ (figure 1, panel D). On the other hand, ⟨2*N*_e_ℙ_fix_⟩ depends weakly on the mutational bias (*λ*).

The proxy of selection represented by ⟨2*N*_e_ℙ_fix_⟩ concerns all non-synonymous mutations, but we can also consider the mean scaled fixation probability only for the subset of non-synonymous mutations from weak nucleotides (A or T) to strong nucleotides (G or C), called 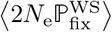. Interestingly, 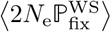 increases with the strength of the mutational bias toward AT (figure 1, panel E). This distortion of the selective effects toward GC is stronger under an increased stringency of selection, under a higher *N*_r_. Likewise, the non-synonymous mutations could also be restricted from strong (GC) to weak nucleotides (AT). This ratio decreases with the strength of the mutational bias toward AT (not shown). As a result, the ratio ratio between 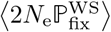 and 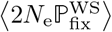 is higher than 1 under a mutational bias toward AT (and lower than 1 respectively for a bias toward GC). It is monotonously increasing with the mutational bias toward AT (figure 1, panel F). Altogether, fixation probabilities are opposed to mutational bias, and the realized equilibrium frequencies are thus at an equilibrium point between these two opposing forces.

### 2.2 Parameter inference on simulated data

From an alignment of protein-coding DNA sequences, without knowing the specific history of substitutions, can one estimate the mutational bias (*λ*) and the mean scaled fixation probability ⟨2*N*_e_ℙ_fix_⟩? In other words, can we tease apart mutation and selection?

To address this question, here we consider two codon models for inference, differing only by their parametrization of the codon matrix ***Q***. Both are homogeneous along the sequence (i.e. not site-specific). The first is based on Muse and Gaut (1994) formalism and uses a scalar *ω* parameter, while the second is based on a tensor representation of *ω*.

#### 2.2.1 *ω* as a scalar: the Muse & Gaut formalism

This model is defined in terms of a generalized time-reversible nucleotide rate matrix ***R*** and a scalar parameter *ω*. The matrix ***R*** is a function of the nucleotide frequencies ***σ*** and the symmetric exchangeability rates ***ρ*** (Tavaré, 1986):

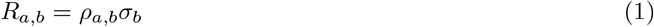

At the level of codons, the substitution rate between the source (*i*) and target codon (*j*) depends on the underlying nucleotide change between the codons ℳ(*i, j*) (e.g. ℳ(*AAT, AAG*) = *TG*), and whether or not the change is non-synonymous. Altogether, the substitution rates between codons *Q*_*i,j*_, formalized by Muse and Gaut (1994) are defined as follows:

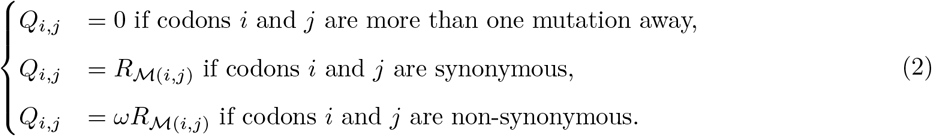

The model can be fitted by maximum likelihood. Then, from the estimate of 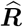, one can derive a nucleotide bias toward AT as:

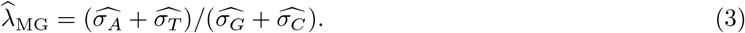

As for the mean strength of selection ⟨2*N*_e_ℙ_fix_⟩, a d irect estimate i s g iven by 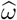.

As shown in the left panel of figure 2, estimate of the mutational bias is halfway between the nucleotide bias observed in the alignment and the true mutational bias used during the simulation. Thus, the MG model cannot reliably infer the mutational bias. On the other hand, 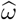 is close to the underlying mean scaled fixation probability ⟨2*N*_e_ℙ_fix_⟩ computed during the simulation (61 primates taxa, 4980 codon sites, 100 repeats), with a precision of 97.2%. Thus, the failure to correctly estimate the mutation process does not seem to have a strong impact on the overall strength selection, at least in the present case.

**Figure 2:**
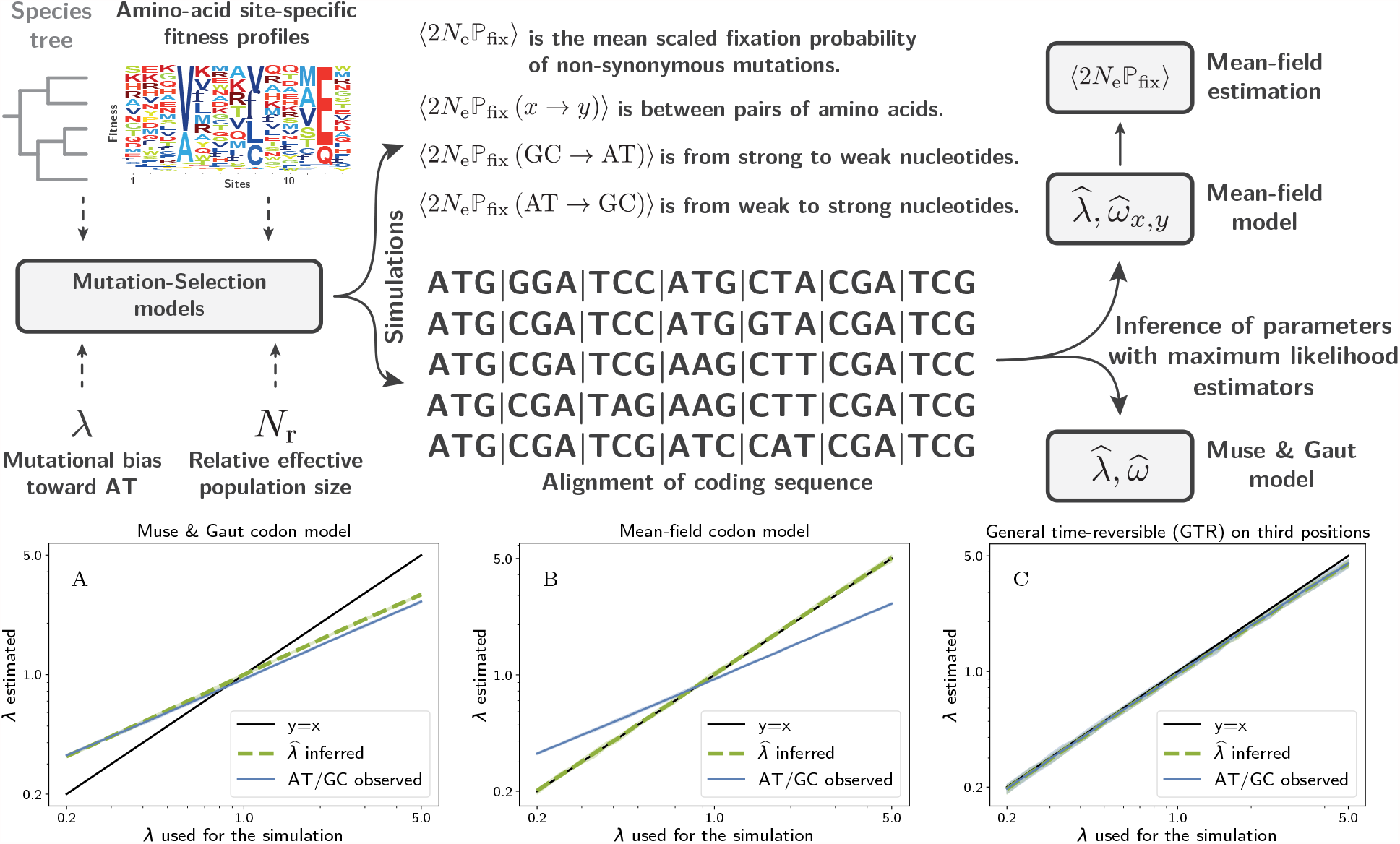
Overall procedure for simulation (61 primates taxa, 4980 codon sites) and inference (top), and estimated versus true mutational bias (bottom), using a codon model in which *ω* is modeled as a scalar (Muse and Gaut formalism, MG, panel A) or as a tensor (mean-field approach, panel B), or by applying a GTR nucleotide model to the 4-fold degenerate third-coding positions only (panel C).

#### 2.2.2 *ω* as a tensor: mean-field derivation

We would like to derive a codon model that would be more accurate than the Muse & Gaut model concerning the estimation of the mutation bias, but that would still be site-homogeneous. However, the true process is site-specific. The link between the two can be formalized by projecting the site-specific processes onto a gene-wise process, using what can be seen as a mean-field approximation (Goldstein and Pollock, 2016). The gene-wise process obtained by this procedure is expressed in terms of mutation rates and mean scaled fixation probabilities. Finally, the mean scaled fixation probabilities can be identified with the *ω*-tensor.

Specifically, at each site z, the true codon process is:

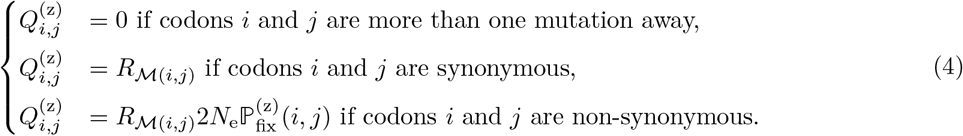

Where 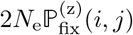 is the scaled fixation probability of codon *j* against codon *i*, at site z. At equilibrium of the process, averaging over sites under the equilibrium distribution gives the mean-field gene-level process:

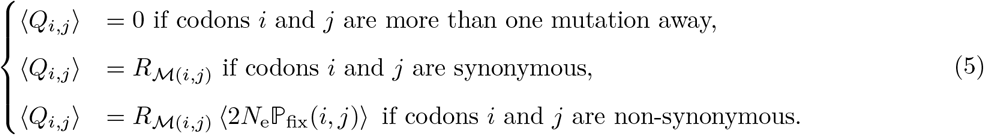

However, because selection between codons reduces to selection between pairs of amino-acids, ⟨2*N*_e_ℙ_fix_(*i, j*)⟩ only depends on the amino-acids encoded by *i* and *j* (section 4.5 in methods). Thus, by identification, the inference model should be parameterized by a set of *ω* values for all pairs of amino acids, denoted *ω*_*x,y*_. For 20 amino acids, the total number of pairs of amino acids is 190, hence 380 parameters by counting in both directions. However, because of the structure of the genetic code, there are 75 pairs that are one nucleotide away, since some amino acids are not directly accessible through a single non-synonymous mutation. As a result, the number of parameters necessary to determine all non-zero entries of the tenser (*ω*_*x,y*_) in both directions is 150. Finally, under the assumption of a reversible process, the number of parameters can be reduced to 75 symmetric exchangeabilities (*β*_*x,y*_) and 20 stationary effects (*ϵ*_*x*_):

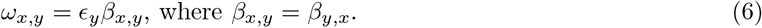

Altogether, the substitution rates between codons *Q*_*i,j*_ are defined as:

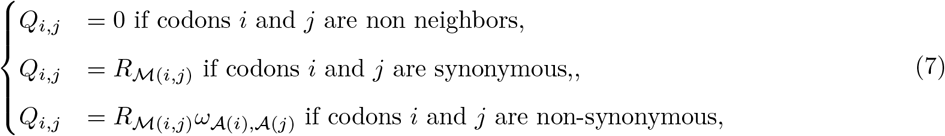

where 𝒜(*i*) is the amino acid encoded by codon *i* and *ω*_*x,y*_ is given by equation 6.

This mean-field (MF) model is fitted by maximum likelihood, giving an estimate for its parameters, 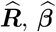 and 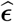. Then, from the estimate of the GTR nucleotide matrix 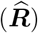, a mutation bias 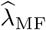 can be estimated as previously (equation 3 above).

As shown in the right panel of figure 2, 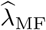 under the MF model provides an accurate estimate of the true mutational. In other words, the MF model can tease out the observed AT/GC bias of the alignment and the underlying mutational bias.

The mean scaled fixation probability of non-synonymous mutations ⟨2*N*_e_ℙ_fix_⟩ can also be computed. It is now a compound parameter, expressed as a function of 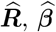 and 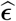 (see section 4.6). Under this model, ⟨2*N*_e_ℙ_fix_⟩ is close to the true mean scaled fixation probability ⟨2*N*_e_ℙ_fix_⟩ computed during the simulation, with a precision of 96.9% (61 primates taxa, 4980 codon sites, 100 repeats). Moreover, as shown in figure 3, the estimated rates 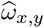 between pairs of amino acids is congruent with the predicted mean scaled fixation probability computed analytically as a function of the underlying site-specific fitness profiles and the mutation matrix as in equation 26.

**Figure 3:**
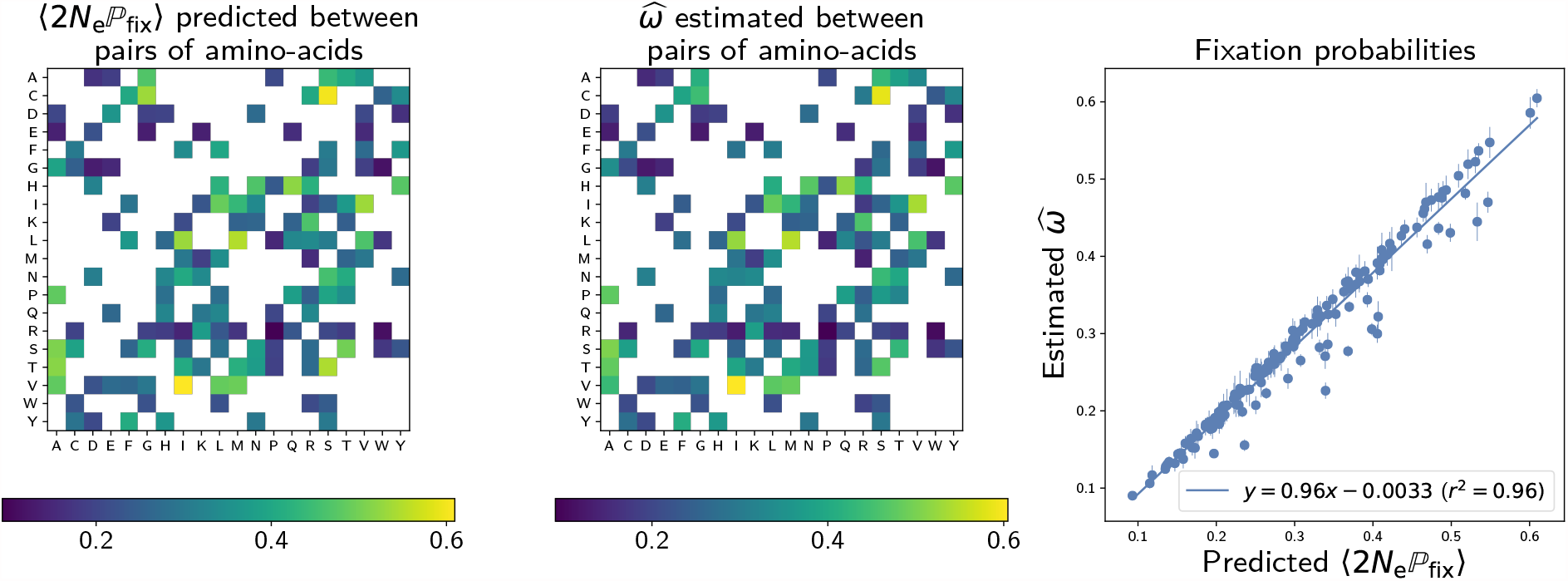
True versus estimated values of *ω* between pairs of amino-acids. The true values are given by equation 7. Simulations on 61 primates taxa with 4980 codon sites over 100 repeats. Vertical bars are the 95% confidence intervals for the mean value.

### 2.3 Estimation on empirical sequence data

The two alternative models of inference just considered, namely the classical Muse & Gaut (MG) and the mean-field (MF) codon models, were then applied to empirical protein-coding sequence alignments. Several examples were analysed: the nucleoprotein in *Influenza Virus* (as human host) assembled in Bloom (2017), the *β*-lactamase in *bacteria* gathered in Bloom (2014), as well as orthologous gene in primates extracted from OrthoMam database (Scornavacca *et al*., 2019) or from Perelman *et al*. (2011) as shown in table 1.

**Table 1:** Mutational bias (*λ*) and mean scaled fixation probability (⟨2*N*_e_P_fix_⟩) estimated under the Muse & Gaut (MG) and mean-field (MF) models on distinct concatenated DNA alignments of orthologous genes.

| | β-Lactamase | Nucleoprotein | Primates AT-rich | Primates |
| --- | --- | --- | --- | --- |
| **Dataset** | **Bloom** | **Bloom** | **Scornavacca *et al.*** | **Perelman *et al.*** |
| **Number of taxa** | 85 | 180 | 22 | 61 |
| **Number of sites** | 263 | 498 | 4877 | 5300 |
| **AT/GC** | 0.792 | 1.154 | 2.028 | 1.075 |
| **AT/GC at 1<sup>st</sup> position** | 0.583 | 1.057 | 1.303 | 0.996 |
| **AT/GC at 2<sup>nd</sup> position** | 1.177 | 1.221 | 2.541 | 1.426 |
| **AT/GC at 3<sup>rd</sup> position** | 0.714 | 1.192 | 2.648 | 0.878 |
| **MG mutational bias ($\hat{\lambda}_{\text{MG}}$)** | 0.853 | 1.447 | 2.073 | 1.139 |
| **MF mutational bias ($\hat{\lambda}_{\text{MF}}$)** | 0.690 | 1.748 | 2.419 | 1.022 |
| **MG $\hat{\omega}$** | 0.332 | 0.114 | 0.526 | 0.272 |
| **MF $\langle 2N_e \mathbb{P}_{\text{fix}} \rangle$** | 0.336 | 0.116 | 0.525 | 0.272 |
| **MF $\langle 2N_e \mathbb{P}_{\text{fix}}^{\text{WS}} \rangle$** | 0.297 | 0.141 | 0.594 | 0.254 |
| **MF $\langle 2N_e \mathbb{P}_{\text{fix}}^{\text{SW}} \rangle$** | 0.412 | 0.092 | 0.487 | 0.308 |
| **$\Delta\text{AIC}$** | 37.6 | 165.2 | 1527.0 | 1091.0 |
| **$p(\chi^2_{\text{df}=93} > \text{LRT})$** | $9.2 \times 10^{-13}$ | $1.2 \times 10^{-31}$ | $3.9 \times 10^{-296}$ | $2.9 \times 10^{-207}$ |

For alignment globally biased toward AT (nucleoprotein and AT-rich concatenate in primates), similarly to what was observed in the simulation experiments presented above, the mutational bias estimates under the two codon models are greater than the observed nucleotide bias (i.e. 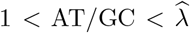). This effect is, as previously, probably due to selection at the level of amino acids, partially opposing the mutational bias. More importantly, the mutational bias estimated by the MF model is more extreme than the MG estimate (i.e. 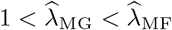). These examples behaves identically to the observations made with simulated alignments, where, compared to MG, the MF model estimates a stronger mutational bias, which was also closer to the real value. Thus, a reasonable interpretation is that MG is also underestimating the underlying mutational bias in the present case, and that the estimate of the MF model is more accurate.

Concerning selection, the estimated mean scaled fixation probability of non-synonymous mutations, is similarly estimated in the MF and MG models 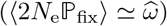. Additionally, in the MF model, ⟨2*N*_e_ℙ_fix_⟩ can be restricted to mutations from weak nucleotides (AT) to strong (GC), or vice versa (see section 4.6). We observe that under a mutational bias favouring AT (i.e. *λ >* 1), the mean fixation probability of non-synonymous mutations is higher toward GC than toward AT, 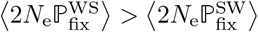, as expected under a AT-biased mutation process.

Reciprocally, for alignment globally biased toward GC (*β*-lactamase), the estimated mutation bias is stronger (toward GC) than the alignment bias (i.e. 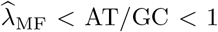). Curiously, in *β*-lactamase, the MG model estimates a weaker underlying mutational bias than the observed bias (i.e. 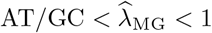).

Concerning selection, we observe that the fixation probability of non-synonymous mutations is higher on average toward AT than toward GC, 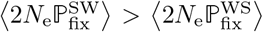, as expected under a GC-biased mutation process.

The results obtained on empirical data are globally in agreement with the observations gathered from the simulation experiments, namely that the presence of a mutational bias results in a selection differential, taking the form of a slightly higher mean fixation probability of non-synonymous mutations opposing the mutational bias. Moreover, by setting ***ϵ***= **1** and ***β*** = *ω* × **1** in our mean-field model, we retrieve the nested Muse & Gaut model, hence, both models are directly comparable. The empirical fit to the data between the nested models, using AIC and Likelihood ratio test (Posada and Buckley, 2004), always favors the MF model compared to the MG model. Altogether, our MF model is favored by empirical dataset, and simultaneously estimates more extreme (and probably more accurate) mutational biases compared to the MG model.

## 3 Discussion

In protein-coding DNA sequences, the nucleic composition results from a subtle interplay between mutation at the nucleic level and selection at the protein level. As a result, the observed nucleotide bias in the alignment is different from the underlying mutational bias.

However, current parametric codon models are inherently misspecified and, for that reason, are unable to tease apart these opposing effects of mutation and selection correctly. As a result, they don’t estimate the mutational process reliably.

In this work we sought to find the simplest parametric codon model able to correctly tease apart mutation rates on one hand, and net mean fixation probabilities on the other hand, and this, without having to explicitly model the underlying fitness landscape. In order to derive a codon model along those lines, our strategy is to first assume an underlying microscopic model of sequence evolution (here, a mutation-selection model based on a site-specific, time-independent fitness landscape). Then, we derive the gene-wise mean fixation probabilities between all pairs of codons, implied by the underlying microscopic process. Finally, we observe that this mean-field process should in fact invoke as many distinct *ω* parameters as there are pairs of amino acids that are nearest neighbours in the genetic code. There are reversibility conditions, reducing the dimensionality and allowing for a GTR-like parameterization of this tensor (95 parameters for selection).

Inferring parameters on simulated alignments, we show that the model derived using this mean-field argument correctly estimates the underlying mutational bias and selective pressure. Applied to empirical alignments, we also observe that there is a selection differential opposing the mutational bias.

This work first points to a fundamental property of natural genetic sequences, namely that they are not optimized but are the result of an equilibrium between forces (Sella and Hirsh, 2005). In the specific case highlighted in this work, mutational bias at the nucleotide-level results in suboptimal amino-acid being overrepresented in the sequence. This was pointed out previously (Singer and Hickey, 2000), although never directly formalized in phylogenetic codon model.

One important consequence of this tradeoff between mutation and selection at equilibrium is that the observed higher mean fixation probability toward GC is mimicking the effect of biased gene conversion toward GC (gBGC), although unlike gBGC, the phenomenon described here corresponds to a genuine selective effect. Although we did not explore the consequences of this at the level of intra-specific polymorphism, the selection differential uncovered here also implies that the distribution of fitness effects is not the same in the two directions, either toward AT or toward GC. Specifically, in the presence of an AT-biased mutation process, the non-synonymous GC polymorphisms are expected to segregate at higher frequencies, compared to non-synonymous AT polymorphisms.

These observations have some practical implications: for instance, experiments observing a fixation (or segregation) bias toward GC at the non-synonymous level must also rule out that this fixation bias is not a simple consequence of the mutation-selection balance. More generally, our observations and modelling principles offer a useful preliminary basis to better understand how mutation and selection will work together with GC-biased gene conversion (gBGC), and therefore will help better understand how gBGC will impact both nucleotide composition and *d*_*N*_ */d*_*S*_. It is worth mentioning that in our result, we focused on the fixation probability from AT to GC, 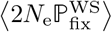, because of the relationship to gBGC. However, in practice, the same analysis and methods can be applied to any subset of nucleotides or codons.

Our mean-field parametric model uses gene-level parameters (in the form of a tensor) that is meant to capture the mean scaled fixation probabilities. This derivation, and its validation on simulated data, shows that, even though the underlying selective landscape is site-specific, a gene-level approximation can nonetheless accurately disentangles mutation and selection. As a result, this study demonstrates that phenomenological models derived out of mechanistic models are more compact (i.e. not site-specific), and in certain cases are sufficient to extract the relevant parameters.

The methodology proposed here for deriving inference models consists in proceeding in two steps, first assuming an underlying mechanistic model of sequence evolution, parameterized by variables that are derived from first principles (fitness landscape, mutations rates, …). Subsequently, the phenomenological inference model is obtained by matching its parameters (here, the entries of the *ω* tensor) with the aggregate parameters derived from the application of the mean-field procedure to the mechanistic model. Altogether, we believe that the approach used here could be applied more generally: inference models can be phenomenological in practice, but should nonetheless be derived from an underlying mechanistic model, so as to correctly formalize the interplay between mutation, selection, drift and other evolutionary forces.

Our phylogenetic codon models is not the first to model *ω* as a tensor, Yang *et al*. (1998) introduced a codon model in which *ω* depends on the distance between amino acids, measured in terms of the Grantham (1974) distance. Additionally, Tang and Wu (2006) leveraged *ω* tensors in order to detect positively selected genes. The novelty of this work is to formalize the articulation between the nucleotide composition, the mutational bias and selection between different amino acids. Finally, this work is still preliminary since the mean-field model should be tested against a more diverse range of empirical data, in terms of phylogenetic depth, strength of selection, and codon usage bias to assert the validity of our empirical results. In addition, several other codon models (Rodrigue *et al*., 2008; Kosakovsky Pond *et al*., 2020) should be included in a broader comparison of the accuracy of the estimation of the underlying mutational bias and strength of selection on protein-coding DNA sequences.

## 4 Materials & Methods

### 4.1 Simulation model

We seek to simulate the evolution of protein-coding sequences along a specie tree. Starting with one sequence at the root of the tree, the sequences evolve independently along the different branches of the tree by point substitutions, until they reach the leaves. At the end of the simulation, we get one sequence for each leaf of the tree, meaning one sequence per species. The substitution is modelled using the origination-fixation approximation, i.e. substitution rates are the product of the mutation rate at the nucleotide level, and fixation probabilities, based on selection at the amino-acid level.

The mutation process is assumed homogeneous across sites. On the other hand, selection is assumed to be varying along the sequence. During the simulation, given the current sequence, the substitution rates toward all possible mutants (one nucleotide change) are computed and the next substitution event is drawn randomly based on Gillespie’s algorithm (Gillespie, 1977).

### 4.2 Mutational bias at the nucleotide level

The mutation rate between nucleotides is always proportional to *µ*. Moreover, mutations from any nucleotide to another weak nucleotide is increased by the factor *λ* compared with mutations to another strong nucleotide.

The mutation rate matrix is thus:

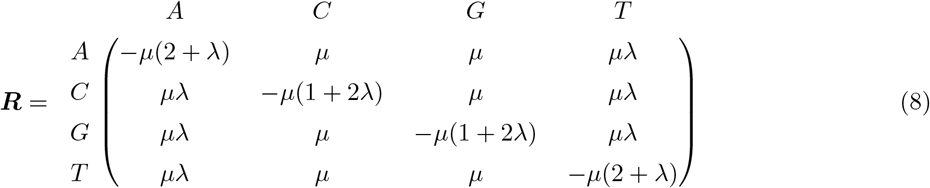

Which has the following stationary distribution:

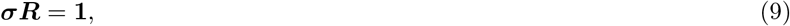

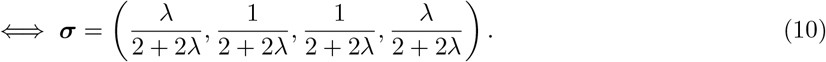

As a result, the ratio of weak over strong nucleotide frequencies at stationarity is equal to *λ*:

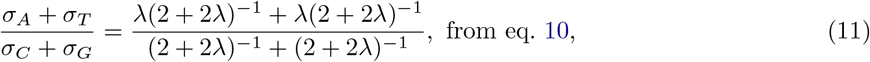

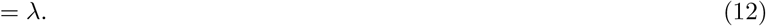

*µ* is constrained such the expected flow (-∑_*a*_ *σ*_*a*_*R*_*a,a*_) of mutation equals to 1.

### 4.3 Selection at the amino-acid level

The substitution rate is considered null between any two codons differing by more than one nucleotide. Otherwise, the mutation rate between a pair of codons is given by the mutation rate of the underlying single nucleotide change. Selection is modelled at the amino-acid level, i.e. we assume that all codons encoding for one particular amino acid are selectively neutral.

To take into account the heterogeneity of selection between different sites of the protein, we assume that each site z of the sequence is independently evolving under a site-specific fitness landscape, characterized by a 20-dimensional frequency vector of scaled (Wrightian) fitness parameters 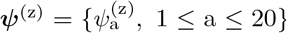. The fitness vectors *ψ*^(z)^ used in this study are extracted from Bloom (2017), which were experimentally determined by deep mutational scanning for 498 codon sites of the nucleoprotein in *Influenza Virus* strains (as human host). For each codon site z of our simulation, we assign randomly one the 498 fitness profile (sampling with replacement) experimentally determined, which altogether determines the (Wrigthian) fitness vectors across sites. The malthusian fitness (or log-fitness) of amino acid a, denoted 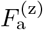, is scaled by the relative effective population size (*N*_r_) accordingly:

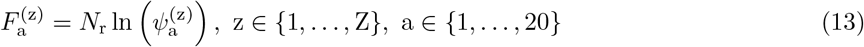

At site z, the substitution rate between non-synonymous codons *i* and *j* is given by the product of the mutation rate and the probability of fixation:

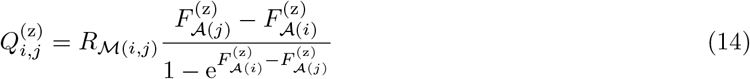

where 𝒜 (*i*) denotes the amino-acid encoded by codon *i*. At the root of the tree, for each site z, the sequence is drawn from the stationary distribution of the process specified by *π*^(z)^, which is given by:

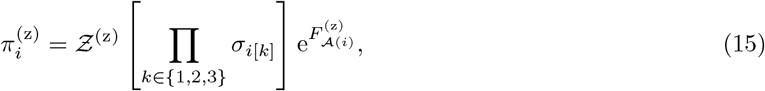

where *i*[*k*] denotes the nucleotide at position *k* ∈ {1, 2, 3} of codon *i*, and Ƶ ^(z)^ is the normalizing constant at site z:

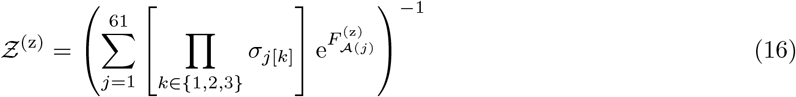

The substitution process is reversible and fulfils detailed balance conditions at each site z and between each pair of codons (*i, j*):

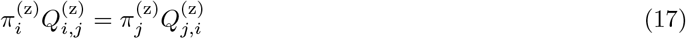

Of note, by modelling fitness at the amino-acid level, we assume that all codons encoding for one particular amino acid are selectively neutral. In addition, in this modelling framework, the genetic code is of particular importance since the number of codons encoding for a particular amino acid varies greatly. As an example, tryptophan is encoded by one codon, while leucine is encoded by 6 codons. Intuitively, this variation makes the mutation bias more pronounced among codons encoding for the same amino acid, since there are more mutations possible that are selectively neutral (i.e. synonymous). On the other hand, the mutation bias is more constrained if the amino acid is encoded by few codons.

### 4.4 Mean scaled fixation probability

The sequence at time *t* is denoted 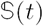 and the codon present at site z is denoted 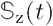. For a given sequence, the mean scaled fixation probability over mutations away from 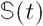, weighted by their probability of occurrence, is given by the ratio:

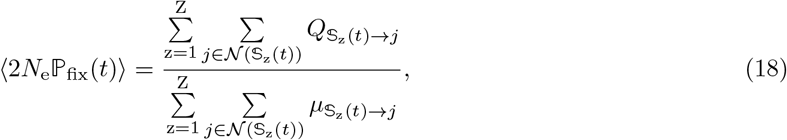

where 𝒩 (*i*) is the set of non-synonymous codons neighbours of codon *i* and 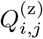 are defined as in equation 14. Averaged over all branches of the tree, the mean scaled fixation probability is :

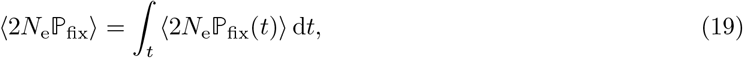

where the integral is taken over all branches of the tree, while the integrand ⟨2*N*_e_ℙ_fix_(*t*) ⟩ is a piece-wise function changing after every point substitution event. The mean scaled fixation probability from weak (AT) to strong (GC) nucleotides, denoted 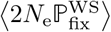, is obtained similarly by restricting the sums (in the numerator and the denominator) from weak to strong mutations. A similar computation can be done from strong to weak.

### 4.5 Derivation of mean-field model

The mean-field codon model ⟨***Q***⟩ is defined such that ⟨*Q*_*i,j*_⟩ is the average rate of substitution to codon *j*, conditional on currently being on codon *i*, the average being taken across sites. Importantly, sites differ in their probability of being currently in state *i*. The average should therefore be weighted by this probability.

Assuming an underlying site-specific mutation-selection process at equilibrium, given we know that a mutation is from codon *i*, the probability that this mutation is occuring at site z is:

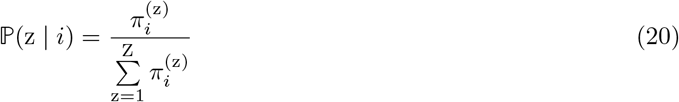

The site-averaged (mean-field) substitution rate from codon *i* to *j* is as result given as:

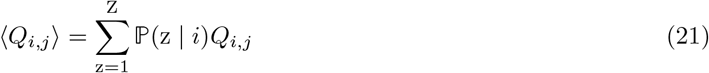

If codon *i* and codon *j* are synonymous, this equation simplifies to the underlying mutation rate *R*_ℳ (*i,j*)_. Otherwise, if codon *i* and codon *j* are non-synonymous, the mean-field substitution rate is:

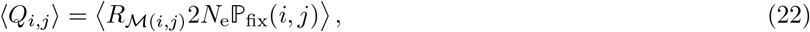

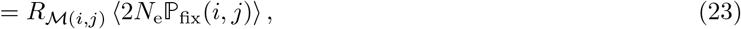

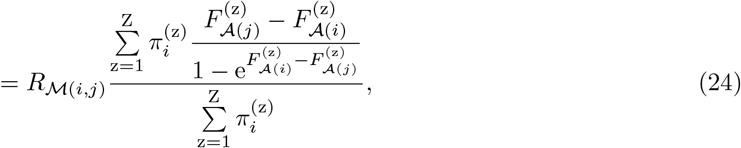

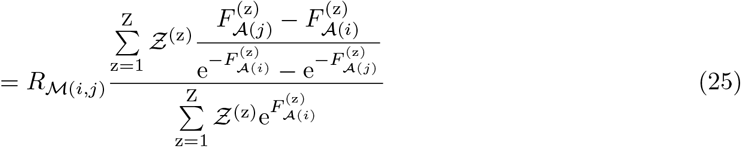

As a result, ⟨2*N*_e_ℙ_fix_(*i, j*) ⟩ is dependent on the source and target codon solely through the source amino acid (*x*) and target amino acid (*y*), hence the parameter *ω*_*x,y*_ identifies with the average fixation probability ⟨2*N*_e_ℙ_fix_ (*x* → *y*)⟩:

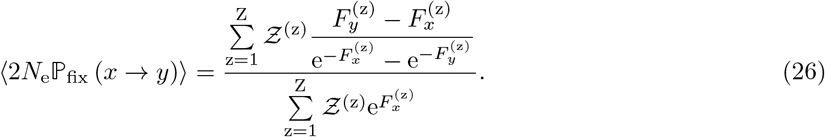

### 4.6 Mean scaled fixation probability ⟨2*N*_e_ℙ_fix_⟩ under the mean-field model

The mean-field model is parameterized by a GTR mutation matrix ***R***(***σ, ρ***) and the selection coefficient ***ω***(***β, ϵ***). As a result, the mean scaled fixation probability of non-synonymous mutations is:

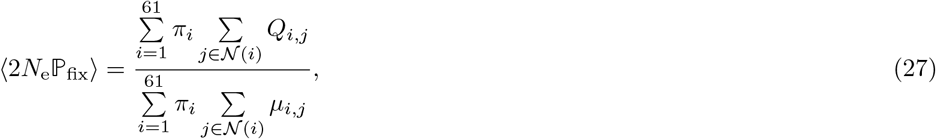

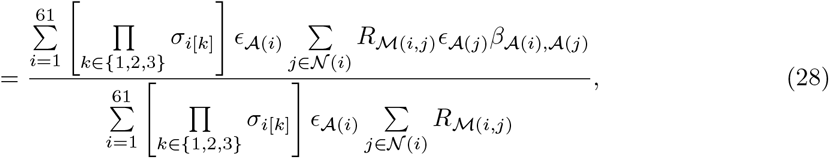

where *i*[*k*] denotes the nucleotide at position *k* ∈ {1, 2, 3} of codon *i*.

Similarly, the mean scaled fixation probability from weak (AT) to strong (GC) nucleotides denoted 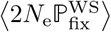 is obtained similarly by restricting the sums (in the numerator and the denominator) to one nucleotide mutations only from weak to strong. Conversely, by restricting the sum from strong (GC) to weak (AT), we obtain 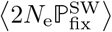.

### 4.7 Inference method with Hyphy

Maximum likelihood estimation has been performed with the software `Hyphy` (Pond and Muse, 2005b). The `Python` scripts generating the `Hyphy` batch files (for both Muse & Gaut and mean-field), as well as scripts necessary to replicate the experiments are available at https://github.com/ThibaultLatrille/NucleotideBias.

## 5 Data availability

The data underlying this article are available in Github, at https://github.com/ThibaultLatrille/NucleotideBias, as well as scripts and instructions necessary to reproduce the simulated and empirical experiments. The simulators written in `C++` are available at https://github.com/ThibaultLatrille/SimuEvol.

## 6 Author contributions

TL gathered and formatted the data, developed the new models in `SimuEvol` and conducted all analyses, in the context of a PhD work (Ecole Normale Superieure de Lyon). TL and NL both contributed to the writing of the manuscript.

## 7 Acknowledgements

We gratefully acknowledge the help of Laurent Gueguen, Laurent Duret, Christophe Douady and Benoit Nahbolz for their input on this work and their comments on the manuscript. This work was performed using the computing facilities of the CC LBBE/PRABI. Funding: French National Research Agency, Grant ANR-15-CE12-0010-01 / DASIRE.

